# No evidence for detectable direct effects of magnetic field on cellular autofluorescence

**DOI:** 10.1101/2022.05.15.491784

**Authors:** Mariia Uzhytchak, Barbora Smolková, Adam Frtús, Alexandr Stupakov, Mariia Lunova, Federica Scollo, Martin Hof, Piotr Jurkiewicz, Gareth John Sullivan, Alexandr Dejneka, Oleg Lunov

## Abstract

Dramatically increased levels of electromagnetic radiation in the environment have raised concerns over the potential health hazards of electromagnetic fields. Various biological effects of magnetic fields have been proposed. Despite decades of intensive research, the molecular mechanisms procuring cellular responses remain largely unknown. The current literature is conflicting with regards to evidence that magnetic fields affect functionality directly at cellular level. Therefore, a search for potential direct cellular effects of magnetic fields represents a cornerstone that may propose an explanation for potential health hazards associated with magnetic fields. Recently, it was postulated that autofluorescence of HeLa cells is magnetic field sensitive, relying on single-cell imaging kinetic measurements. Here, we explore the utility of this approach by undertaking a screen for magnetic sensitivity of cellular autofluorescence in statistically relevant numbers (90-107) of HeLa cells. We did not observe any changes in cellular autofluorescence decay, when a modulated magnetic field was applied. We present a number of arguments indicating weak points in the analysis of magnetic field effects based on the imaging of cellular autofluorescence decay. Our work indicates that new methods are required to elucidate the effects of magnetic fields at the cellular level.

## 1. Introduction

Despite decades of research, the biological effects of magnetic fields still remain a highly debatable topic without consensus over major outcomes [1-4]. Thus far, dozens of studies ascribed a bewildering variety of biological effects to both electromagnetic as well as static magnetic fields [5-8]. It has been widely criticized that many investigations which deal with the biological impact of magnetic fields are hampered by shortcomings in experimental design and often also by the lack of reproducibility [1-4, 6, 7, 9, 10]. Specifically, direct attempts to replicate key findings on biological effects of magnetic fields were largely unsuccessful [11-15]. Therefore, it is not surprising that the molecular and/or biophysical foundations for the alleged cellular effects still remain enigmatic.

Furthermore, some epidemiological studies presumed a low interrelation between residential proximity to high-voltage power lines and childhood leukemia [16-18]. However, recent thorough literature analysis showed that majority of large-scale epidemiological studies do not support this association (for review see [19] and references therein). In this flow of contradictory data the International Agency for Research on Cancer (IARC) classified magnetic fields of extremely low-frequencies as “possibly carcinogenic to humans” (Group 2B) in 2002, despite “*inadequate*” and “*limited*” evidence [20]. In 2011, the IARC once again reviewed studies on health risks of magnetic fields and concluded that the evidence was inadequate to clearly link magnetic field exposure with adverse health impacts, leaving magnetic fields in Group 2B of carcinogenicity to humans [21, 22]. Recent systematic reviews on bioeffects of weak and intermediate, static and different frequency range, magnetic and electromagnetic fields found, that current evidence does not allow to draw a firm conclusion for biological and health-related consequences of exposure to those fields [4, 23, 24]. Therefore, it is very premature to claim that current research has identified harmful influence of magnetic fields on human health [25].

It is worth noting, to directly establish the existence of a health impact of a certain agent, it is crucial to have solid evidence from epidemiological analysis, empirical laboratory animal studies and *in vitro* studies of the agent’s cellular mechanism of action [19]. Magnetic fields-associated adverse health effects derive primarily from disputable epidemiological studies [19]. Of note, *in vivo* animal and *in vitro* mechanistic studies suggest a lack of support for health risks resulted from magnetic field exposure [19]. Thus, research on revealing direct potential mechanisms of magnetic fields at the cellular level is extremely interesting and a hot topic. Deciphering cellular targets may propel our understanding and support our understanding of magnetic field health impacts. In fact, numerous studies on biological effects of magnetic fields have produced many claims, proposing distinct potential mechanisms of magnetic field action on biological matter [1-3, 6-8]. One tentative hypothesis explaining biological effects of magnetic fields is alteration of biochemical reactions via so-called radical pair (RP) mechanism (RPM) [3, 26, 27]. Indeed, attempts to independently reproduce suggested magnetic field effects (MFEs) via RPM in biochemical systems were unsuccessful [11, 28-32]. It is worth noting here, that there are studies that doubted the relevance of RPM hypothesis in biological systems and questioned the interpretation of results [2, 33, 34]. It was proposed that flavin adenine dinucleotide (FAD)-containing photoreceptors (cryptochromes) might be a key player responsible for RPM-based MFEs [35-37]. However, others showed that reactions catalyzed by flavin-dependent enzymes are unlikely to be influenced by magnetic fields [38]. So far, studies evaluated the hypothesis of flavin-based RPM in cell-free systems [37-39]. The recently published study proposed, that there might be magnetic field sensitivity of FAD-based RPM at the cellular level [25]. In the light of all discussed above, MFEs at the cellular level represent an extraordinary claim. We absolutely agree that, in order to deliberately show biological relevance of MFEs, it is an imperative to ***robustly*** and ***directly*** demonstrate magnetic field responses at individual cell level [25], because *extraordinary claims require extraordinary proof*. Therefore, we were very keen to reevaluate the recently published findings of Ikeya and Woodward, that showed MFE at cellular level [25]. Here we evaluate MFE on the autofluorescence of HeLa cells, introducing several critical control points that were omitted in the original study.

## 2. Materials and methods

### 2.1. Cell culture

HeLa cells were obtained from the American Type Culture Collection (ATCC CCL-2). Cells were cultured in Minimum Essential Medium Eagle (BioConcept Ltd., Switzerland) supplemented with 10 % fetal bovine serum (FBS, Thermo Fisher Scientific, US), 1% L-Glutamine 100X, 200 mM (Serana Europe GmbH, Germany) and 1 % Penicillin/ Streptomycin (Thermo Fisher Scientific, US). Cell cultures were cultivated in a humidified 5% CO_2_ atmosphere at 37 °C. Cell culture medium was replaced once a week.

### 2.2. Magnetic field exposure setup

Magnetic field was generated by a pancake/bobbin coil of 3 cm width and 11/14 cm inner/outer diameters having 550 turns of copper wire of 0.8-mm diameter. The coil was calibrated using a laboratory Gaussmeter FW Bell 7030 and their transverse Hall probe STF71-0404-05-T with temperature compensation. The coil was adjusted directly on the top of Olympus IX3-SVR mechanical stage. Cells were placed directly inside the coil, in the central axial area on a level of the coil edge. Specific number of the coil in this area was estimated as ∼5 mT/A. Triangular voltage waveforms with frequencies of 0.1-0.2 Hz were supplied by a standard arbitrary waveform generator OWON AG1022 (Owon, China). The driving signal was power amplified in a voltage control mode by a MP39 module mounted on an EK59 evaluation kit (APEX Microtechnology, US). During the measurements, the magnetic field was controlled by the same Gaussmeter and transverse Hall probe positioned at the opposite upper coil edge. Amplitude of the magnetic field was set to 10 or 20 mT. Driving current ∼4 A (33 V) was applied to generate 20 mT magnetic field. DC offset of the magnetic field measured without driving current does not exceed 0.1 mT and mostly determined by the Earth’s magnetic field.

### 2.3. Autofluorescence decay of HeLa cells upon exposure to magnetic field

Cell were seeded in 6-channels Ibidi μ-slides (Ibidi, Germany) and propagated until 80-85% of confluence. After, cell culture medium was replaced with preheated calcium/magnesium free PBS buffer. Cells were washed twice with PBS buffer before the live imaging.

We performed autofluorescence magnetic field measurements in a very similar way to the methodology described in [25]. We focused on HeLa cells utilizing bright-field illumination to avoid photobleaching. Afterwards, cells were imaged under continuous irradiation with 2.95 mW of 488-nm laser light and an applied magnetic field varying between either +10 mT and –10 mT or +20 mT and –20 mT at frequencies of 0.1 Hz or 0.2 Hz. The laser power was measured with an optical power and energy meter PM100D (Thorlabs Inc., US) using S170C microscope slide power sensor (Thorlabs Inc., US). Digital built-in thermometer inside the generator was used to monitor absence of heating effect of samples during magnetic field exposure. The irradiation intensity was estimated to be 0.4 kW/cm^2^ on the sample. Fluorescent images were captured with a camera exposure time of 100 ms.

In order to treat cells with static magnetic field, we applied cylindrical (radius 5 mm; length 50 mm; residual magnetic flux density 1,4 T) bulk NdFeB magnet. Magnet was applied on the top of the 6-channels Ibidi μ-slides (Ibidi, Germany) at the central part of the channel (Figure S1). Magnetic flux density (*B(x)*) at the level of cells was estimated to be ∼ 500 mT (Figure S1).

In order to cross-check that the noise from imaging system and other equipment in the laboratory does not have significant impact, we measured the background electromagnetic noise. To estimate this the voltage induced in the same bobbin coil was measured by a 16-bit acquisition board NI PCIe-6351 at 2 MSa/s sampling rate. The measured noise voltage of ∼8 mV amplitude and 2.5 mV rms value is independent of the microscope activity (whether it is switched on/off) and fully determined by background electromagnetic noises in the laboratory (Figure S2A). RMS value of the highest noise harmonic at 639 kHz is ∼1.5 mV; the highest harmonic in a low-frequency range is of ∼0.2 mV rms at power line frequency 50 Hz (Figure S2B). Such a 50-Hz harmonic gives a leap of the magnetic field ∼0.15 µT, which is a common level of the background magnetic noise.

### 2.4. High-resolution fluorescent imaging

In order to get high-quality fluorescent images for further autofluorescence decay analysis, the high-resolution spinning disk confocal system IXplore SpinSR (Olympus, Japan) was used. The system utilizes an inverted microscope (IX83; Olympus, Japan) and a spinning disc confocal unit (CSUW1-T2S SD; Yokogawa, Japan). Fluorescence images were obtained through 100x silicone immersion objective (UPLSAPO100XS NA 1.35 WD 0.2 silicone lens, Olympus, Tokyo, Japan). Autofluorescence was excited by 488 nm laser. Confocal images were acquired at a definition of 2,048 × 2,048 pixels. A bandpass filter (BA510-550; Olympus, Japan) was used before scientific Complementary Metal Oxide Semiconductor (sCMOS) camera ORCA-Flash4.0 V3 (Hamamatsu, Japan). Images were taken with the acquisition software cellSens (Olympus, Japan).

### 2.5. Image processing and data analysis

Fluorescent signal from a single cell (defined by ROI) was defined as the sum of pixel intensity for a single image with the subtracted average signal per pixel for a region selected as the background. Such analysis was done using function *intensity profile* in software CellSens (Olympus, Japan). We analyzed autofluorescence decay of 90 to 109 individual cells per condition. We performed three independent experiments on different days. To compare level of an integrated density, we used ImageJ software (NIH, US).

Curve fitting and residual analysis were done in SigmaPlot 13.0 software (Systat Software Inc., US). Global curve fitting was conducted using a single exponential decay function (*f = y0+a*exp(-b*x)*). Normalized residuals were calculated as (obtained value − fitted curve value)/(fitted curve value). Additionally, we calculated MFE, defined as *[I(B*_*0*_*) – I(0)]/I(0)*, where *I(B*_*0*_) and *I(0)* are the fluorescence/absorption intensities in the presence and absence of the magnetic field respectively [37, 39, 40].

### 2.6. Fluorescent probes

To highlight differences in brightness of synthetic fluorescent probes and endogenous fluorescence, we labelled cells with standard fluorescent dyes. Cells were labeled with CellMask™ Green (Thermo Fisher Scientific, US) in order to visualize plasma membrane. Additionally, mitochondria were stained with MitoTracker® Green FM (Thermo Fisher Scientific, US). Stained cells were imaged using the spinning disk confocal microscope IXplore SpinSR (Olympus, Japan).

### 2.7. Fluorescence spectra measurements

The fluorescence spectra of HeLa cells and a solution of Atto488 were measured with a FS5 spectrofluorometer (Edinburgh Instruments Ltd., UK). The excitation was performed with a Xenon-Arc Lamp light with a selected excitation wavelength of 450 nm. The excitation slits and the corresponding light powers were 1 and 5 nm, and 0.06 and 1.2 mW, for the synthetic dyes and for the autofluorescence of the cells, respectively. The fluorescence was acquired from 480 nm to 800 nm using 1 nm resolution and a dwell time of 0.2 s with 3 times averaging.

### 2.8. Statistical analysis

The sample size determination was assessed utilizing a statistical method described in [41], taking into assumption 95% confidence level and 0.9 statistical power. The statistical significance of differences between the groups was determined using ANOVA with subsequent application of Dunnett’s test. All statistical analyses were performed using MaxStat Pro 3.6. Differences were considered statistically significant at (*) *p* < 0.05.

For a quantitative image assessment, we used the published guidance for quantitative confocal microscopy [42, 43]. Images from three independent experiments were subjected to quantitative analysis. In each experiment, at least 90 cells from each sample were subjected to quantitative analysis.

## 3. Results

### 3.1. Autofluorescence of HeLa cells and quantitative fluorescence microscopy

Concept of MFE at cellular level, proposed in [25], is based on the hypothesis that an external magnetic field affects the photochemistry of flavins. Specifically, it can influence the spin-correlated radical pairs formed via intersystem crossing in the excited state of a flavin molecule. This would change the rate of flavin deexcitation and the concentration of flavin molecules in the ground state. As a result, a change in the cellular endogenous fluorescent signal should be observed under continuous photoexcitation [25]. In other words, the autofluorescence photobleaching should be altered by external magnetic field. Furthermore, the study states that flavins are the major source of cellular autofluorescence in the range 480-650 nm when excited with blue laser (450 nm) [25].

Healthy undamaged cells possess relatively low level of autofluorescence in comparison to modern synthetic exogenous fluorophores [44-46], which are nowadays greatly optimized to offer higher quantum yield and brightness (Table S1). In fact, HeLa cells imaged in phosphate-buffered saline (PBS) showed relatively dim autofluorescence (Figures 1A and S3). We utilized very similar conditions for excitation (“blue” 488 nm laser) and emission detection (BA510-550 nm “green” filter and sCMOS camera ORCA-Flash4.0 V3) as in the published study [25]. However, we made a few modifications, which were crucial in our opinion, for a more thorough and robust analysis of the autofluorescence signal. In the original study [25], only a part of a cell was irradiated by a laser spot. We irradiated all cells captured in the field of view to increase the sampling size. Actually, electromagnets used in the original study and in ours are relatively bulky, which precludes focusing of the magnetic field on a single cell. Thus, since all cells on the microscopy slide are subjected to the magnetic field, it is logical to illuminate all of them in order to get more sampling for a better statistical assessment of the results. In the original study [25], it is noted that cells showed a magnetic response with a magnitude of 1 to 2.5%. These are very weak changes. Guidance for quantitative fluorescence microscopy state that for a change of even as high as 25% between two conditions, sampling of ∼ 100 cells for each condition is required to measure the change with statistical confidence [42]. In fact, in the study of Ikeya and Woodward [25] it is stated that “*a large number of individual HeLa cells for different cell cultures prepared on different days*” was used to analyze autofluorescence. Unfortunately, we could not find a definition of “*a large number*”, nor any other sample size determination. The largest number of analyzed cells was indicated as 11 in Supplementary Figure S1 [25]. Another essential change we introduced, when analyzing the data for autofluorescence decay, was a background correction. It is absolutely necessary for accuracy and precision in quantitative fluorescence microscopy measurements, especially when quantifying weak signals [42, 47].

**Figure 1.**
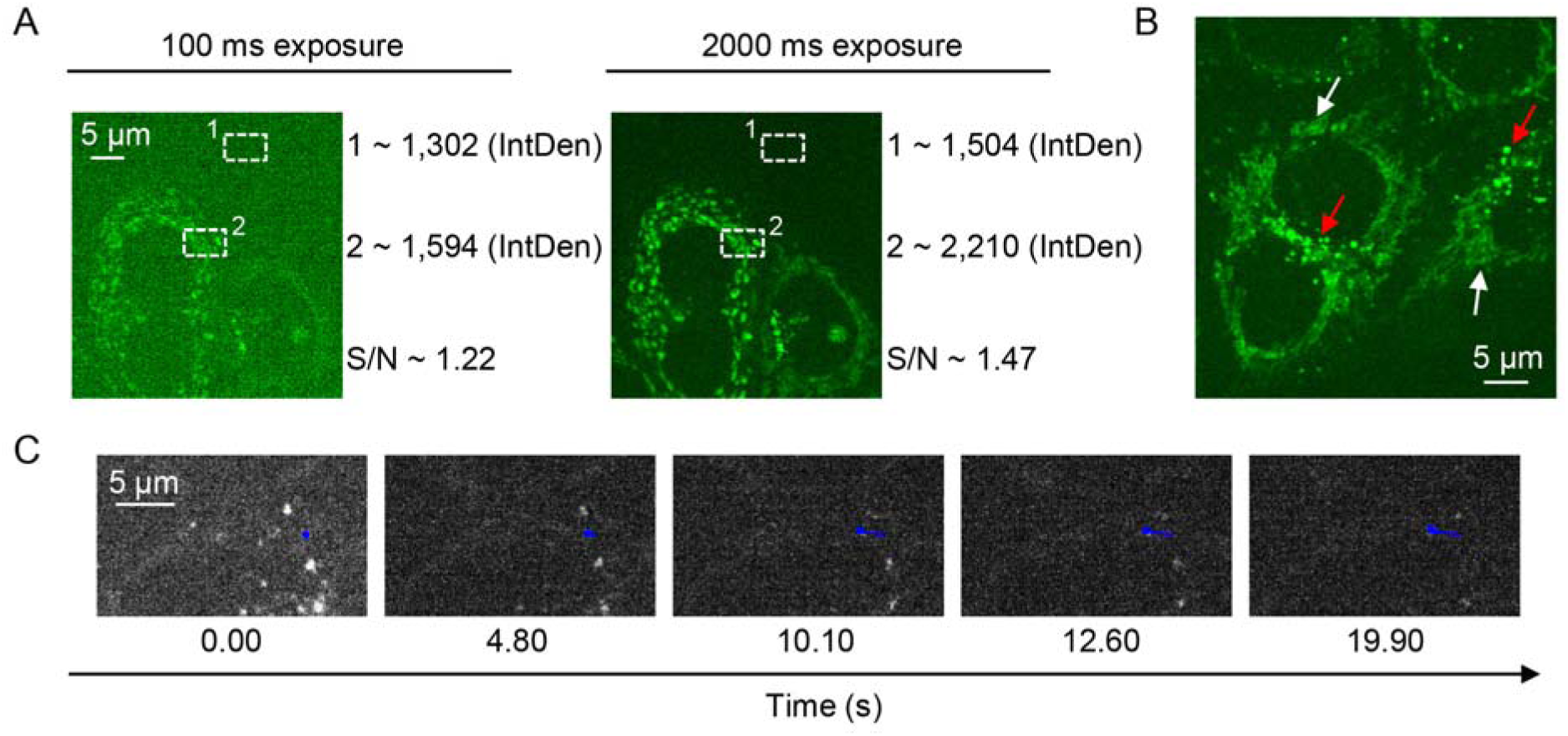
Autoflourescence of HeLa cells. (**A**) Autoflourescence of HeLa cells at 100 ms and 2000 ms exposure time. Integrated density (*IntDen*) was measured for the background (1) and cell region (2) using ImageJ software (NIH, US). Signal-to-noise ratio (*S/N*) was calculated for image selections. (**B**) Enlarged picture represents tubular (white arrows) and round (red arrows) intracellular structures. 2000 ms exposure time. (**C**) Vesicle tracking within the time range, 0-20 sec. Blue color represents vesicle movement through the time-lapse image (Movie S3).

Further analysis of HeLa cell autofluorescence revealed that one can get a better imaging using a longer (2 s) exposure time (Figures 1A and S3). This is typical in detection of dim fluorescent signals [42, 47]. Interestingly, there are various endogenous fluorophores that contribute to “green” (500-550 nm) autofluorescence of cells [44, 46], for details see Supporting Information, Table S2. Not only flavins are responsible for such autofluorescence, but also lipofuscin and free fatty acids (Table S2). Figure 1B and Supporting Information, Figure S4 show that “green” autofluorescence originates from at least two distinct subcellular structures, i.e. tubular and vesicular. The main subcellular structures contributing to “green” autofluorescence are, indeed, mitochondria and lysosomes [48, 49]. Autofluorescence from mitochondria was proposed to come from flavins (for example, FAD), whereas lysosomal autofluorescence is presumably due to lipofuscin accumulation [48, 49]. Taking together those studies and our results (Figure 1B), we can say that it is not single subcellular structure contributing to the autofluorescence. The fact that autofluorescence of HeLa cells is weak (Figure 1A) is important for quantitative analysis of the images, due to Poisson noise persistence in fluorescence microscopy digital images [47]. For example, bright synthetic exogenous fluorophores provide better suitable images (Figure S5) for quantitative analysis in contrast to autofluorescence (Figure 1A).

From the captured autofluorescence photobleaching movie (Movie S1) it is clear that the autofluorescent structures are moving. To present moving structures more clearly, we zoomed several cells (Movie S2). Additionally, we performed particle tracking analysis of selected vesicles (Figure 1C and Movie S3), and found them moving approx. 2.5 μm in 20 seconds (Figure 1C and Movie S3). Weak fluorescence signal in combination with fast-moving structures add disturbance in further quantitative image analysis [42, 47]. Moreover, different cells display different levels of autofluorescence intensity (Figure S6). As a result, autofluorescence decay upon photobleaching in different cells largely varies (Figure S7 and Movies S4 and S5). Further, some cells contain both highly movable and non-movable fluorescent entities (Figure S7 and Movie S6).

As one can see, in the light of all above mentioned, measurements of “green” autofluorescence decay are highly variable and possess huge number of uncertainties.

### 3.2. MFE measurements in HeLa cells

In the Ikeya and Woodward study [25], different amplitudes (e. g. ± 40, ± 30, ± 25, ± 20 and ± 10 mT) and frequencies (between 0.05 and 0.25 Hz) of magnetic field were applied perpendicular to the sample cover glass with living cells. We intended to conceptually verify MFE on HeLa cells. Thus, we selected amplitudes (± 20 and ± 10 mT) and frequencies (0.1 and 0.2 Hz) of magnetic field. In the original study, they did not perform sample size determination nor any reasonable statistical analysis [25]. It is critical to estimate the minimum number of replicates needed for multilevel regression to avoid biases. Importantly, only large effects can be detected using small sample size [50]. A small sample size was shown to result in overestimates of effects and low reproducibility of results [51]. In fact, the study [25] reveals that “*cells showed a magnetic response with a typical magnitude of 1 to 2*.*5 %*”. This is a small effect, requiring appropriate sampling estimation. For such a small effect, it is important to have at least 100 replicates in order to get sufficient statistical power needed for obtaining significance [52]. Therefore, in our study we analyzed autofluorescence decay of 90-109 individual cells per condition (Figures S8 and S9). However, only performing replicates is not enough; repeating experiments should be conducted in different days [53]. Therefore, we performed three independent experiments on different days (Figures S8 and S9).

One can see that autofluorescence decay (Figure S8) is very variable within a cell population, even in controls (without magnetic field). Some cells, even without applied field, show fluctuating “spikes” of autofluorescence during photobleaching course (Figures S8 and S9). In order to clearly examine these changes in autofluorescence, we sub-selected from 12 to 18 of the most “responding” cells (Figure 2A). We added one more control, that is a static magnetic field (SMF), generated by a bulk magnet. SMF was applied perpendicular to the sample cover glass. In fact, one can see, that “spikes” of autofluorescence occur both with and without application of external field (Figure 2A). The pattern is consistent even after averaging of the autofluorescence change (Figure 2B). Neither amplitude nor frequency of the “spikes” of autofluorescence are dependent on the applied field amplitude and frequency (Figure 2). In addition, when one looks closely to individual cell autofluorescence change (Figures S8 and S9), one can see that there is no pattern changed upon application of magnetic field of different amplitude and frequency. The outcome gets even clearer, when averaging of 90-105 cells (Figures 3A and S10). It is very hard to assume the presence of any effect with such decays of autofluorescence (Figures 3A and S10). Plotted on the same graph averaged autofluorescence intensity curves look very similar (Figure 3A).

**Figure 2.**
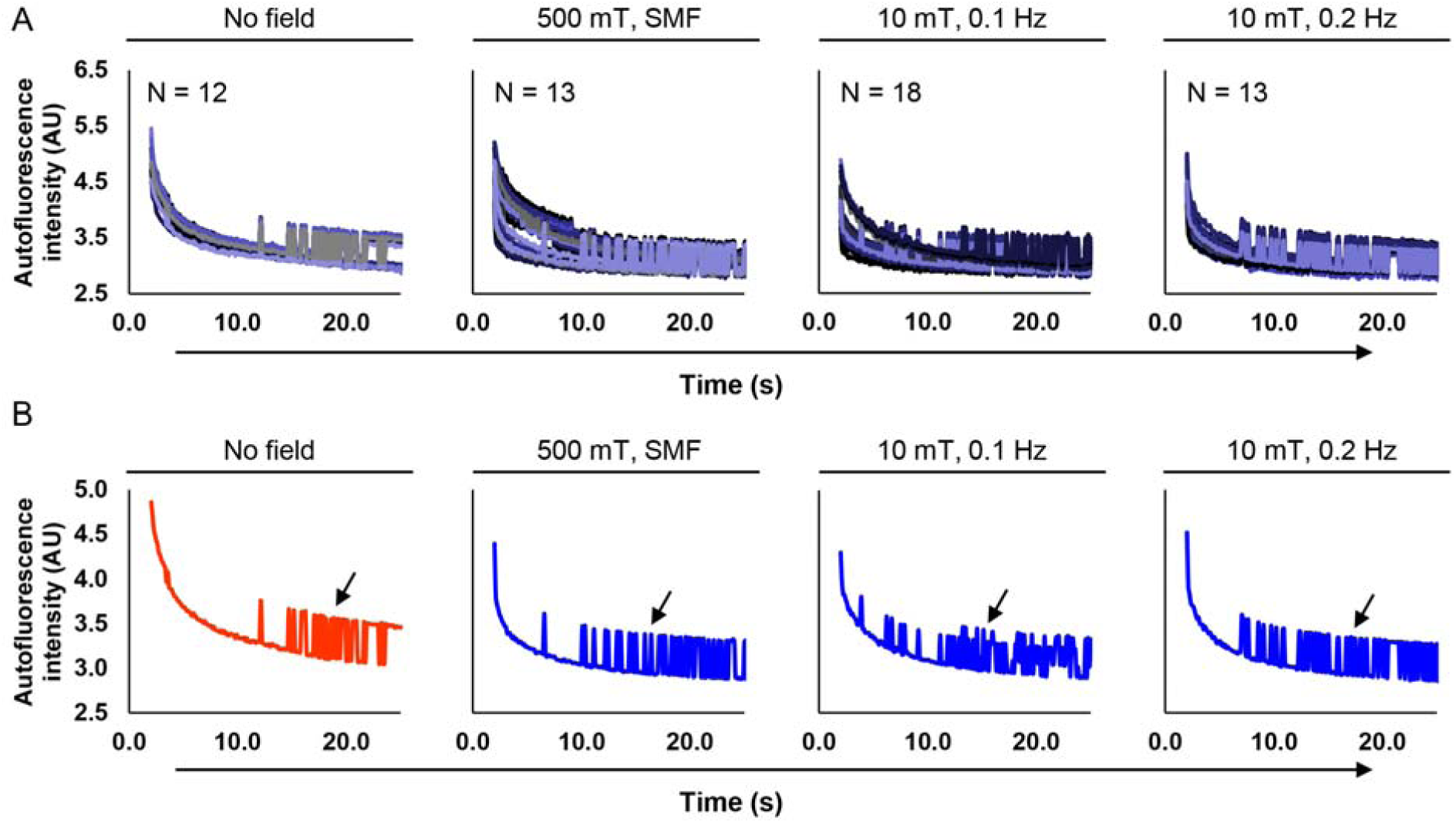
Autoflourescence decay of selected HeLa cells upon a magnetic field exposure. (**A**) Autofluorescence intensity within time-frame of 20 sec for selected number of cells for control (without magnetic field exposure), bulk magnet (500 mT) exposure and a modulated magnetic field 10 mT (frequencies 0.1 Hz and 0.2 Hz). (**B**) Averaged autoflourescence decay of cells presented in (**A**). Arrows indicate oscillating patterns.

**Figure 3.**
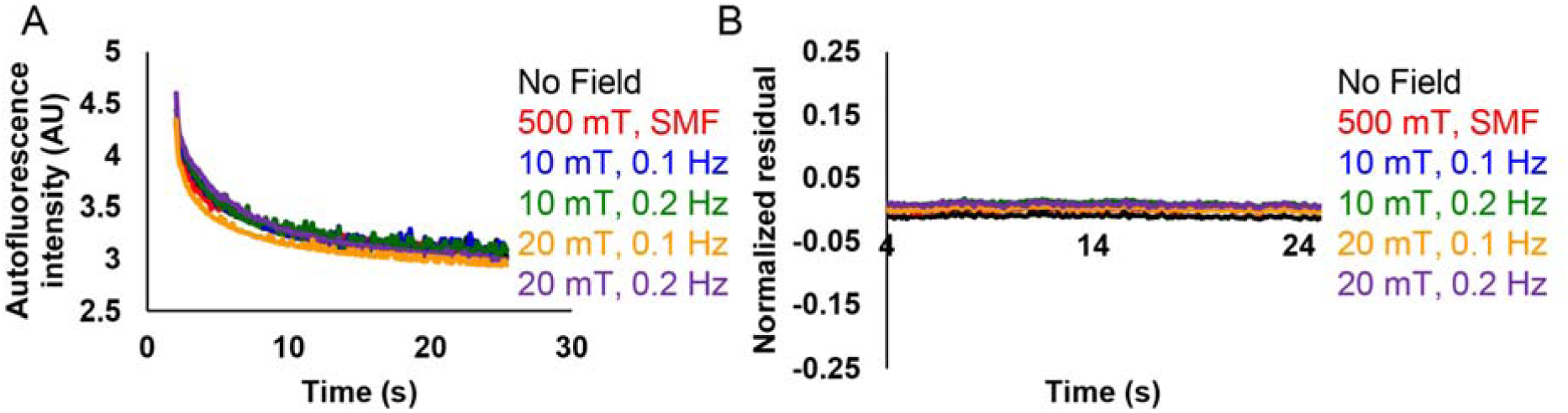
Averaged autofluorescence intensity change of HeLa cells. (**A**) Average autofluorescence decay upon different magnetic field exposure. Cells were irradiated by 10 mT and 20 mT modulated magnetic field (frequencies 0.1 Hz and 0.2 Hz). 500 mT static magnetic field (SMF) was generated by bulk NdFeB magnet. N=90-107. (**B**) Normalized residuals calculated from the average autofluorescence intensity divided by the obtained intensity from the fitted curve values within 20 sec. N=90-107.

### 3.3. Gambling with statistics and mathematical analysis

We were puzzled by how the fractional MFE was estimated in the Ikeya and Woodward study [25]. The average autofluorescence decay curves were fitted using *some* exponential function [25]. We could not define; what kind of exponential function exactly was used. In the methods section “*double exponential function*” was stated, whereas in the main text and in the supplements “*single exponential function*” were mentioned [25]. Indeed, averaged autofluorescence decay curves can be nicely fitted used either single exponential or double exponential 5 parameters decay functions (Figure S11). Double exponential decay 4 parameters function showed a bad approximation of the data, reflected by low values of R-square and Adjusted R-square, as well as high error value (Figure S11). For further analysis, we utilized the most frequently mentioned in the text of the original study “*single exponential function*”. Then normalized residuals were calculated as (observed value − fitted curve value)/(fitted curve value) [25]. These residuals were postulated as a “*measurement of the fractional MFE*” [25]. Different kind of residuals may be used in regression analysis. However, residuals, generally, are used to detect various types of disagreement between data and the assumed model [54]. Basically, residual analysis shows the quality of the regression. The residuals randomly distributed around zero indicating the validity of a particular selected regression model for a given data set [54]. Contrary, fluctuating patterns of residuals around zero over time would suggest that the error term is dependent [54]. In other words, it is a prerequisite of the uncertainty in the model, that strongly suggests a lack of perfect goodness of a fit [54].

Indeed, calculated normalized residuals for 90-107 cells showed a random distribution around zero (Figures S12 and S13). We were unable to observe any consistent patterned changes in the residuals corresponding to the magnetic field frequency or amplitude (Figures S12 and S13). Corresponding averaged values of normalized residuals only support absence of any pattern associated with the MFE (Figures 3B, S12 and S13). Interestingly, we could sub-select a small number of cells in control (without any field exposure), which showed either fluctuating or non-fluctuating patterns of residuals around zero (Figure S14). To clarify, some HeLa cells randomly without any magnetic field exposure exhibited clear fluctuating patterns of residuals (Figure S14).

However, we think that the major shortcoming of this study, that have led to the misinterpretation of the results [25], was making inference without directly comparing two effects (e.g. autofluorescence in the presence and in the absence of magnetic field). In fact, this is very common mistake leading to inappropriate analysis in the literature [50]. Making conclusion about an effect of some treatment should be based on a direct statistical comparison between a control and a treatment group [50]. We directly compared levels of cell autofluorescence at the 20th second after photobleaching in the presence and absence of magnetic fields (Figures 4A and S15). We analyzed levels of cell autofluorescence of 90-105 individual cells. One can clearly see, that there is no statistically significant difference in the autofluorescence between the control cells (no magnetic field) and the cells exposed to any of the magnetic field conditions used (Figure 4A).

**Figure 4.**
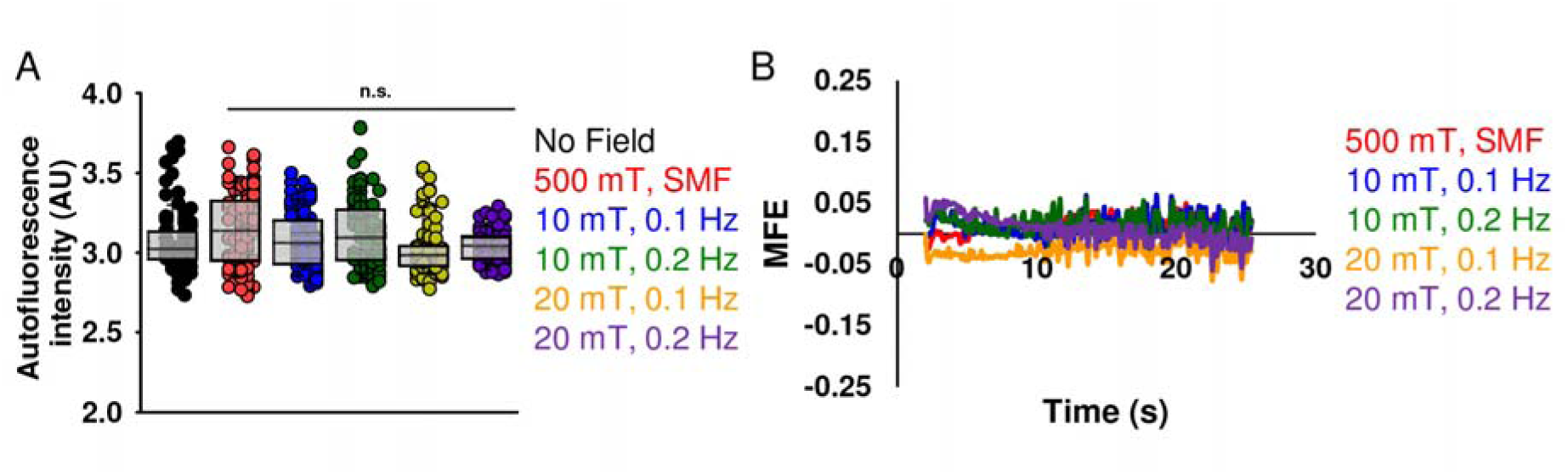
Magnetic field effect on HeLa cells autoflourescence intensity (**A**) Autofluorescence intensity comparison measured 20 sec after exposure to different magnetic fields. Cells were irradiated by 10 mT and 20 mT modulated magnetic field (frequencies 0.1 Hz and 0.2 Hz). 500 mT static magnetic field (SMF) was generated by bulk NdFeB magnet. N=90-107. Dunnett’s test was used to determine statistical significance. Differences were considered statistically significant at *P < 0.05. (**B**) Magnetic field effect (MFE) in the presence of static and a modulated magnetic field for averaged autofluorescence data calculated from Fig. 3A.

It is worth noting here that in the previous works dealing with MFEs on chemical reactions, MFE was defined as *[I(B*_*0*_*) – I(0)]/I(0)* [37, 40] or *[I(0) – I(B*_*0*_*)]/I(0)* [39]. *I(B*_*0*_) and *I(0)* are the fluorescence/absorption intensities in the presence and in the absence of the magnetic field respectively [37, 39, 40]. We did further calculations of MFE, defined as *[I(B*_*0*_*) – I(0)]/I(0)*. It is very hard to assume any trend in such disordered data (Figure 4B). Neither frequency nor amplitude dependence of MFE were observed in HeLa cells upon magnetic field exposure (Figure 4B).

In order to stress that autofluorescent signal from cells is weak; we measured fluorescence spectra in HeLa cells (Figure 5). However, one can see that fluorescent signal from cells is orders of magnitude lower in comparison with standard Atto488 dye (Figure 5A). Moreover, spectrum from cells had significant impact from Raman scattering of water (Figure 5B). Generally, fluorescence can be more intense than the weak Raman scatter, and Raman interference is not presenting an issue for fluorescence spectra measurements. In fact, in diluted solutions of fluorophores the Raman scatter from the solvent can significantly distort the measured fluorescence spectrum [55]. This is what we have observed with HeLa cells (Figure 5B). Interestingly, even after the solvent spectrum subtraction and normalization the HeLa cells spectrum had multiple peaks (Figure 5C), not as smooth as normalized Atto488 spectrum (Figure 5D). These findings only support our conclusions that autofluorescence of HeLa cells is weak and supposedly comprises of distinct chemical entities (e.g. FAD, FMN, lipofuscine, glycation adducts see Table S2).

**Figure 5.**
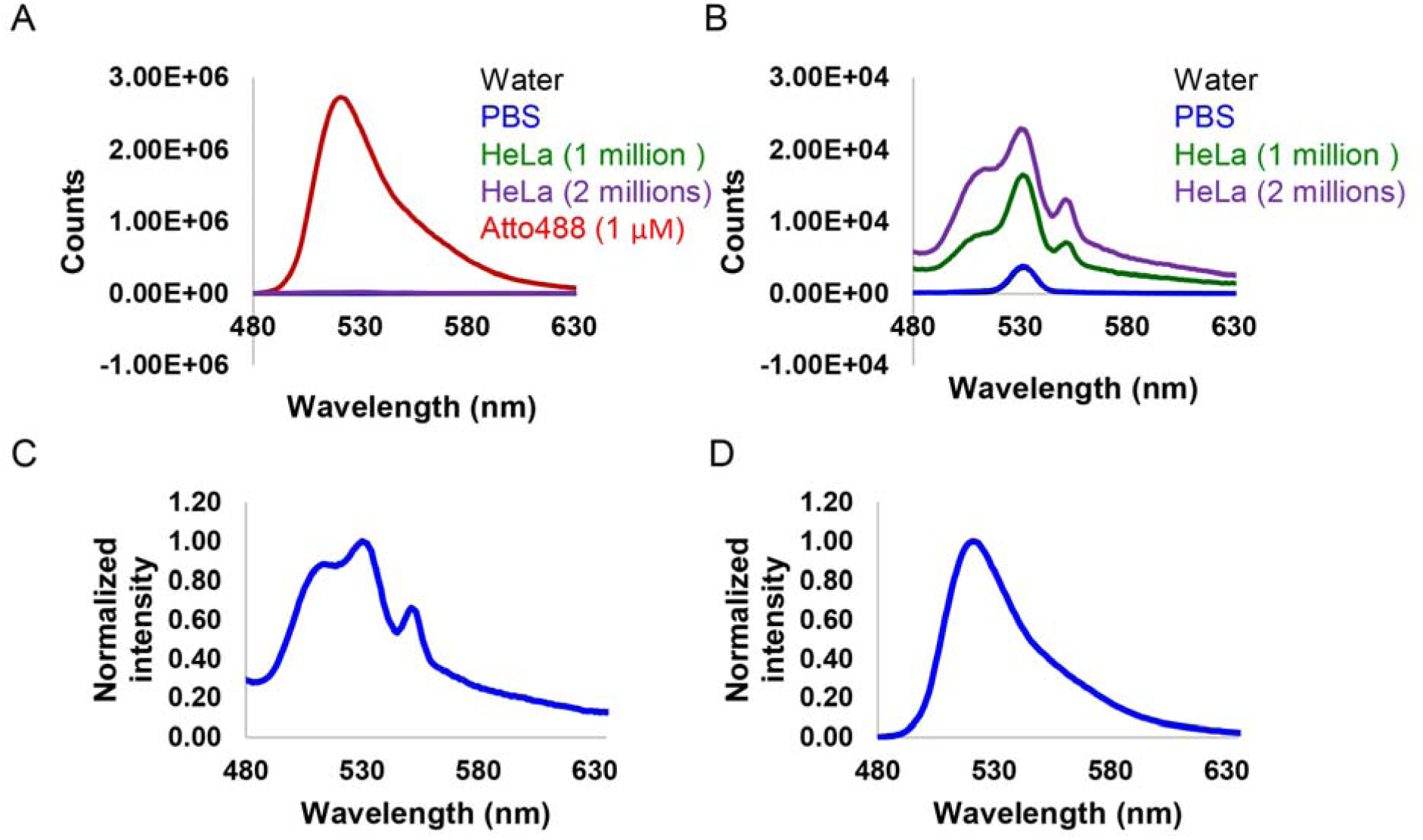
Fluorescence spectra of HeLa cells under 450 nm excitation. (**A**) Fluorescence spectra of water, PBS, HeLa cells diluted in PBS and Atto488 (1 μM in PBS). (**B**) Enlarged fluorescence spectra of water, PBS, HeLa cells diluted in PBS. (**C**) Normalized to the maximum intensity and adjusted for the Raman scattering fluorescence spectrum of HeLa cells diluted in PBS. (**D**) Normalized to the maximum intensity fluorescence spectrum of Atto488 (1 μM in PBS).

## 4. Discussion

Direct identification of biological effects of magnetic fields still remains elusive [1-4]. A lack of reproducibility of key findings is still a major challenge in this field of research [1-4, 6, 7, 9, 10]. When systematic or critical reviewing is applied to analyze inconsistencies in reports of the biological effects of magnetic fields, it quite often leads to the conclusion, that there are either insufficient/erroneous descriptions of design, execution, or validation of the experimental methods and systems [2, 4, 6, 7, 10]. Detailed reporting of the key elements of experimental setup, as well as, mathematical and statistical analysis of the generated data are crucial for the effective reproducibility of the results and establishing findings as scientifically verified facts [56]. Neglecting these guidelines may lead to problems and the claimed findings may only represent measures of the prevailing bias [57]. Due to the irreproducibility of the results on biological effects of magnetic fields [11-15], it is important to verify independently claims about MFEs, especially at a cellular level. The multidisciplinary area of biological effects of magnetic fields suffers from dramatic variability in experimental details reported, usage and characterization of biological models. This, in turn, creates a significant barrier to progress research forward. Therefore, we suggest that research community should establish a ‘minimum information standard’ for research dealing with biological effects of magnetic fields, such as already existing for biochemical models [58], genome sequencing [59], quantitative PCR [60], animal research [61]and bio–nano experimental literature [62].

In this study, we reexamined the claims, that “*endogenous autofluorescence in HeLa cells is sensitive to the application of external magnetic fields*” [25]. Based on cellular autofluorescence decay measurements, we show that this is not the case and the observed autofluorescence is not magnetic field dependent and appears to be an artifact of their analysis. We show this as follows, we focused on the sources of endogenous autofluorescence. In fact, there are many endogenous compounds responsible for 500-550 nm range of autofluorescence (Table S2). Flavins are only a part of the cocktail of compounds contributing to cellular “green” autofluorescence (Table S2). We demonstrate, that “green” autofluorescence is highlighted by at least two (i.e. tubular and vesicular) distinct subcellular (compartments) structures (Figures 1B and S4). Our findings are in line with literature that identifies the mitochondria and lysosomes as the main subcellular structures that contribute to endogenous autofluorescence [48, 49]. Of note, lysosomal autofluorescence is a result of lipofuscin accumulation [48, 49]. Thus, it is very premature to neglect the contribution of lipofuscin in cellular autofluorescence signal. Especially, when original studies used a longpass filter (ET500lp, Chroma) [25]. One will certainly detect both “yellow” and “red” range of fluorescent signal utilizing such a filter. In such a case, the resulting autofluorescence will be a sum of signals from NAD(P)H, flavins, lipofuscins, retinoids, porphyrins, bilirubin and lipids [44, 46]. Although we used more specific bandpass filter (BA510-550), two different sub-cellular structures were still detected (Figures 1B and S4). Of note, detection systems (i.e. sCMOS camera ORCA-Flash4.0 V3) were identical. Levels of autofluorescence intensity vary between different cells (Figure S6). Autofluorescent structures can move at relatively high speed (Figure 1C and Movie S3). All these factors contribute to the large variability in the observed autofluorescence decay upon photobleaching (Figure S7 and Movies S4 and S5), and this observed variability will spill over into the quantitative analysis of images [42, 47]. Indeed, this is not surprising, because the level of cellular autofluorescence depends on multiple factors, e.g. metabolic activity [63], cell cycle phase [64], cell aging and level of oxidative stress [65], degree of cell damage and cell death [66, 67].

Given complexity of endogenous sources of autofluorescence and multiple factors affecting its levels, it is of no surprise, that the origin of autofluorescence photobleaching is still not completely understood [68]. As a result, several distinct decay models can be used for fitting the photobleaching dynamics, ranging from one-to three exponential decay functions [68-70]. Thus, it is not understandable what criteria were used in the selection of fitting model in the study of Ikeya and Woodward [25]. We have shown that photobleaching dynamics can be nicely fitted using two different models (Figure S11).

Additional confusion arises on the introduction of “*measurement of the fractional MFE*” [25], calculated as normalized residuals. The residuals are defined as (observed value − fitted curve value)/(fitted curve value) [25]. In the absence of direct evidence of MFE at the cellular level, such calculation can only show the goodness of the selected fitting model [54]. Direct comparison of control cells with cells exposed to a magnetic field is requireed to make a reasonable interpretation. It is very nicely summarized in [50], how omitting direct comparison can lead to flawed conclusions. This is exemplified in our findings that the application of a magnetic field of different amplitude and frequency did not result in any noticeable effect on the autofluorescence decays in HeLa cells (Figures 3A and S10).

Another oversight, made in the study of Ikeya and Woodward [25], is the use of small sampling size i.e. number of cells analyzed. The “*typical*” magnitude of a magnetic response was reported from 1 to 2.5 % [25]. With such small size effects one should be extremely cautious about interpretation and thus acquire appropriate sampling numbers to validate their observation [50]. Utilization of small sample sizes precludes low reproducibility of results and potentially exaggeration of the observed effect [51]. When we acquired a large enough sampling, the previously observed patterned changes in the residuals corresponding to the magnetic field frequency or amplitude were not detected (Figures S12 and S13). The averaged values of normalized residuals underline absence of the MFE in HeLa cells (Figures 3B, S12 and S13). Initial huge turbulence in the autofluorescence decay signal, multiple endogenous and exogenous factors affecting its levels, spurious calculations of “*the fractional MFE*”, all may have led to observations of fluctuating patterns of residuals around zero importantly in a small number of cells (sampling), even in the absence of field exposure (Figure S14). It must be noted, that our experimental approach, in some respects, differs slightly from that described by Ikeya and Woodward [25] (see Materials and Methods). However, none of these minor differences would be expected to significantly affect the proposed mechanism of “*the fractional MFE*” in HeLa cells.

Shortcomings in the experimental design and methodology, compounded by a lack of reproducibility are the factors highlighted in studies about the biological impact of magnetic fields [1-4, 6, 7, 9, 10]. Additionally, even though some epidemiology studies speculate on tentative relation between the exposure to magnetic fields and health-related effects, independent verifications of the main results show no clear evidence [4, 19, 23, 24]. Therefore, it is very important to critically analyze and importantly independently corroborate statements about MFEs in living cells.

In conclusion, we were not able to reproduce the key results postulated in the study of Ikeya and Woodward [25]. More importantly, we highlight a number of arguments revealing the reasons for their interpretation. Collectively, we conclude that the approach proposed in the study of Ikeya and Woodward [25] was not suitable to reveal the MFE at a cellular level, due to (i) turbulent and weak autofluorescent signal from single cells, (ii) dependence of the autofluorescence on multiple factors, (iii) cellular autofluorescence representing a sum of signals contributing from distinct endogenous compounds and (iv) small sample size and low statistical power of the study.

## Appendix A. Supplementary data

Supplementary data to this article can be found online at https://doi.org/

### Abbreviations

RP: Radical pair
RPM: Radical pair mechanism
MEFs: Magnetic field effects
FAD: Flavin adenine dinucleotide
SMF: Static magnetic field

## Author Contributions

O.L. designed research; M.U., B.S., A.F., A.S., M.L., F.S. performed research; O.L., M.U., B.S., A.F., A.S., M.L., F.S., P.J. M.H., and A.D. analyzed data; O. L. wrote the original draft; M.U., B.S., A.F., A.S., M.L., P.J., M.H., G.J.S., and A.D. reviewed and edited the paper; and O.L. conceived the project.

## Availability of data and material

All the data in the current research are available from the corresponding author on reasonable request.

