## Supplementary data for "No evidence for detectable direct effects of magnetic field on cellular autofluorescence"

Number of pages: 18

Number of tables: 2

Number of figures: 15

Number of movies: 6

**Table of Contents**


| | |
|---|---|
| Table S1 | S3 |
| Table S2 | S3 |
| Figure S1 | S4 |
| Figure S2 | S4 |
| Figure S3 | S5 |
| Figure S4 | S5 |
| Figure S5 | S6 |
| Figure S6 | S7 |
| Figure S7 | S7 |
| Figure S8 | S8 |
| Figure S9 | S9 |
| Figure S10 | S10 |
| Figure S11 | S11 |
| Figure S12 | S12 |
| Figure S13 | S13 |
| Figure S14 | S14 |
| Figure S15 | S15 |
| Legends for Movies | S16 |
| References | S17 |

**Table S1.** Quantum yield and brightness of different compounds.

| Molecule | Quantum yield | Molar extinction coefficient ($M^{-1}$ $cm^{-1}$) | Brightness[1] | Ref. |
|---|---|---|---|---|
| Riboflavin | 0.36 | 33000 | 11880 | (1, 2) |
| Flavin mononucleotide (FMN) | 0.26 | 12200 | 3172 | (3, 4) |
| Flavin adenine dinucleotide (FAD) | 0.033 | 11300 | 372.9 | (5, 6) |
| Tryptophan | 0.12 | 5540 | 664.8 | (7, 8) |
| Fluorescein (FITC) | 0.95 | 76000 | 72200 | (9, 10) |
| Alexa Fluor 488 | 0.92 | 73000 | 67160 | (11) |

[1]The brightness is a product of extinction coefficient and quantum yield of the fluorophore (12).

**Table S2.** Selected endogenous fluorophores responsible for cell and tissue autofluorescence.

| Molecule | Excitation, peak position range (nm) | Fluorescence, peak position range (nm) | Ref. |
|---|---|---|---|
| Flavin adenine dinucleotide (FAD) | ~ 380-490 | ~ 520-560 | (13) |
| Flavin mononucleotide (FMN) | ~ 380-490 | ~ 520-560 | (13) |
| Lipofuscin | ~ 410-488 | ~ 500-695 | (14) |
| Elastin | ~ 350-420 | ~ 420-510 | (15) |
| Glycation adducts of collagen | ~ 370-420 | ~ 450–460 | (16) |
| Free fatty acids (arachidonic, linoleic, linolenic acid) | ~ 330-350 | ~ 470-480 | (17) |
| Biliary salts and bilirubin | ~ 400-490 | ~ 540-600 | (17) |

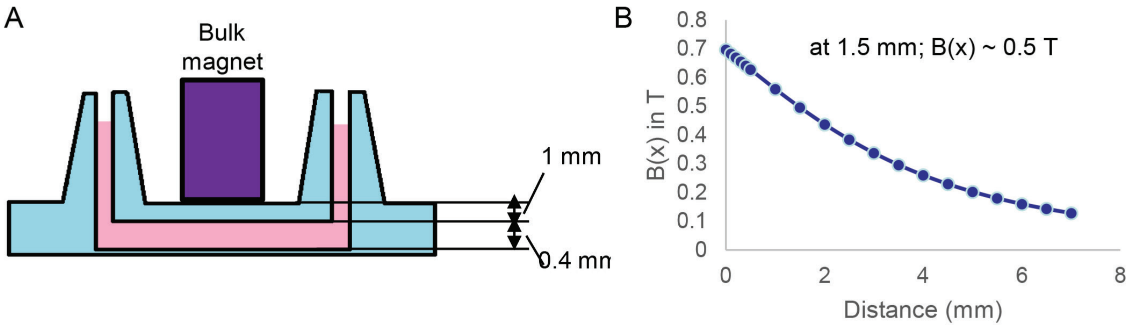


**Figure S1.** Estimation of the magnetic field generated by bulk NdFeB magnet. (*A*) Schematic application of bulk NdFeB magnet. (*B*) Calculation of distance decay of the flux density B(x), utilizing approached described in (18).

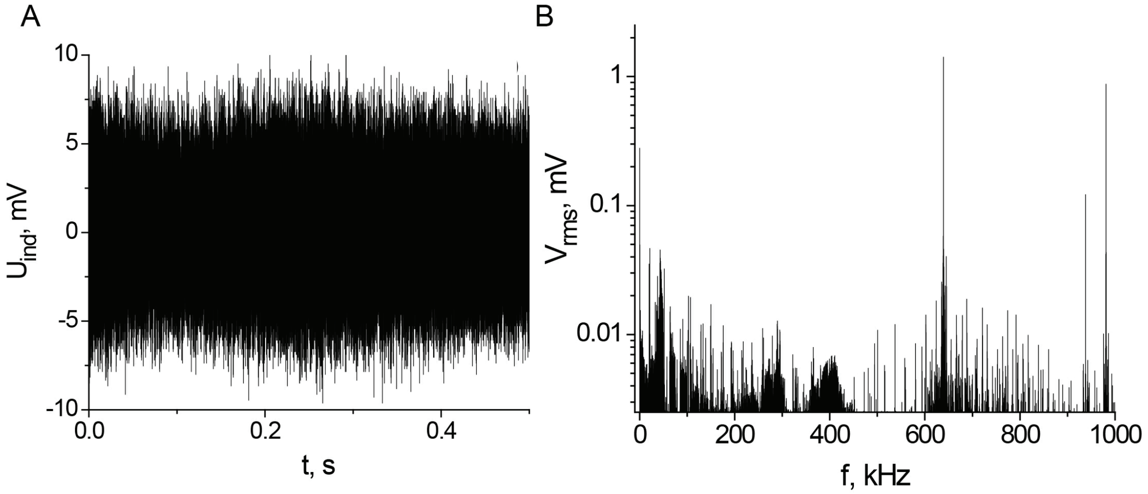


**Figure S2.** Measurements of magnetic environmental noise. (*A*) Background electromagnetic noise measurements. (*B*) Fourier power spectrum of the background electromagnetic noise.

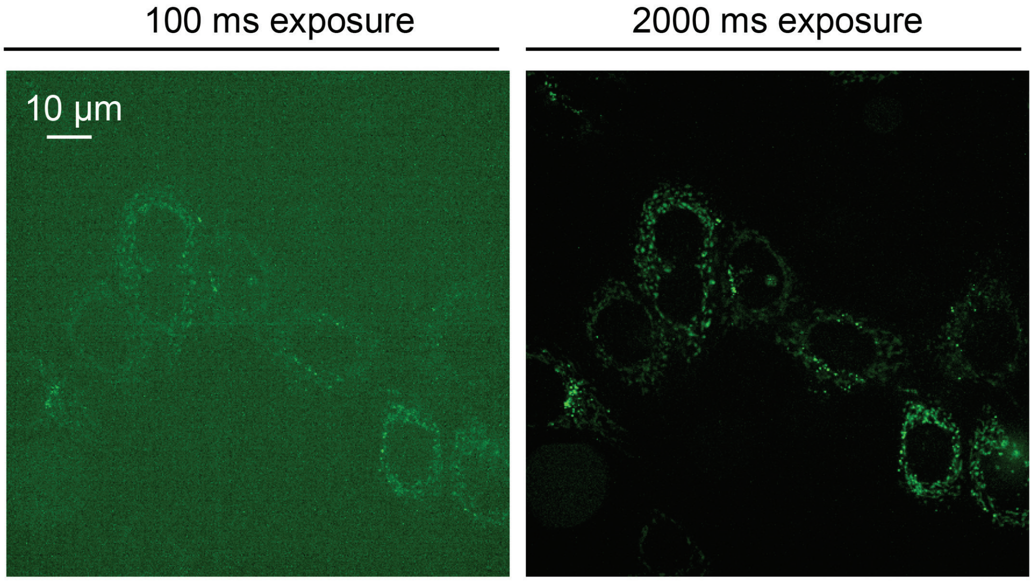


**Figure S3.** Full view of HeLa cells autoflourescence confocal images. Autoflourescence images of HeLa cells taken with 100 ms and 2000 ms exposure time.

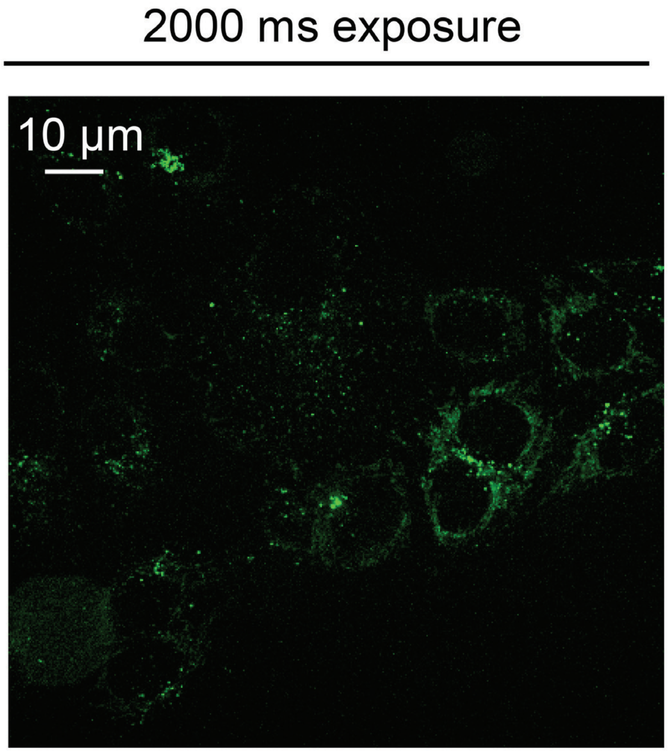


**Figure S4.** Full view image representing tubular and round intracellular structures in HeLa cells taken with 2000 ms exposure time.

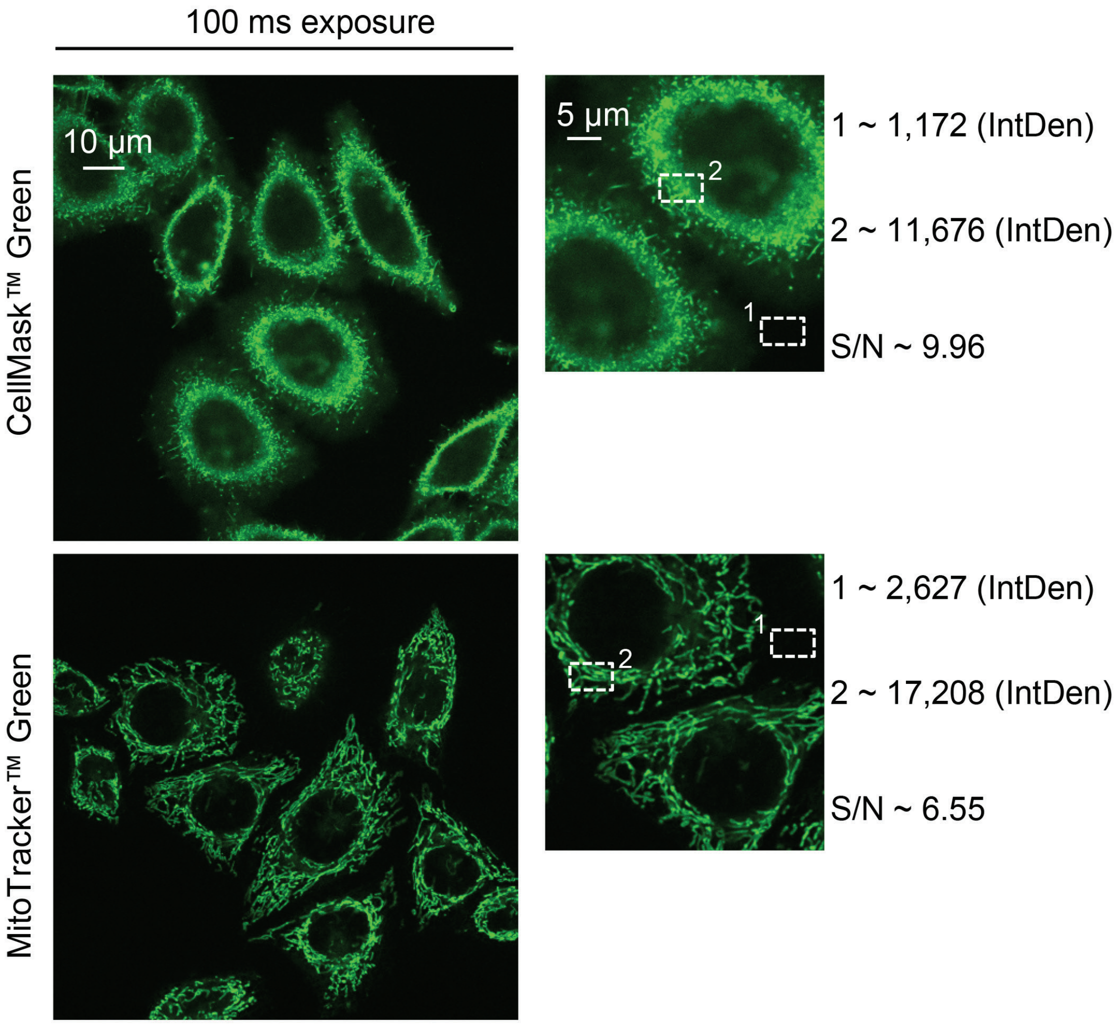


**Figure S5.** Fluorescent intensity measurements of HeLa cells stained with standard probes. Cells were labeled with CellMask™ Green (Thermo Fisher Scientific, US) in order to visualize plasma membrane. MitoTracker® Green FM (0.5 µM), (Thermo Fisher Scientific, US) was used for mitochondria staining. Signal-to-noise ratio (S/N) is calculated for selected image regions. Integrated density (*IntDen*) was measured for the background (1) and cell part (2) using ImageJ software (NIH, US).

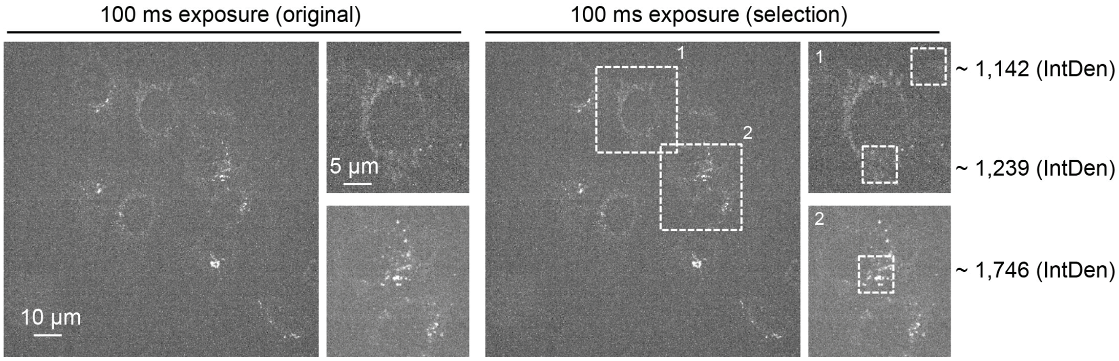


**Figure S6.** Cell autoflourescence presented in a grayscale images with 100 ms exposure time. Integrated density (*IntDen*) was measured for the background and cells regions using ImageJ software (NIH, US).

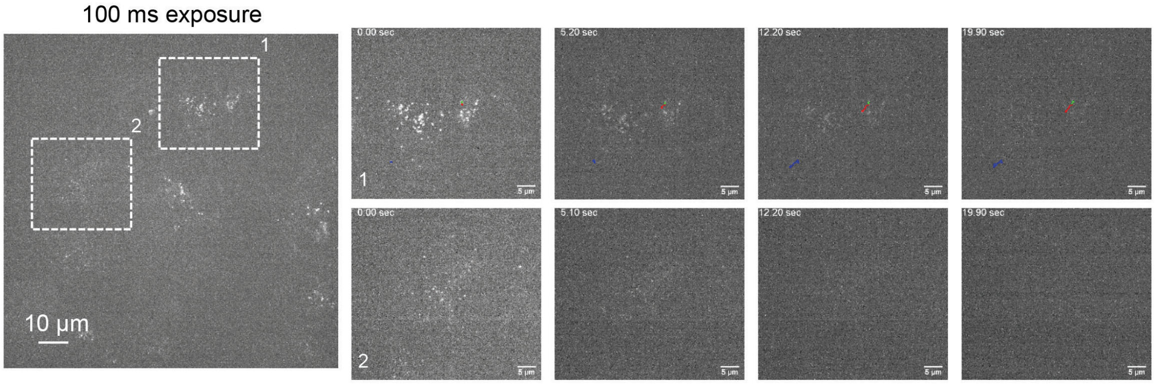


**Figure S7.** Autoflourescence decay time-lapse of HeLa cells within 20 sec period of laser irradiation. Cell autoflourescence dacay presented in a grayscale images with 100 ms exposure. Blue, red and green colors represent vesicle movements through the time-lapse image (Movies S4, S5 and S6).

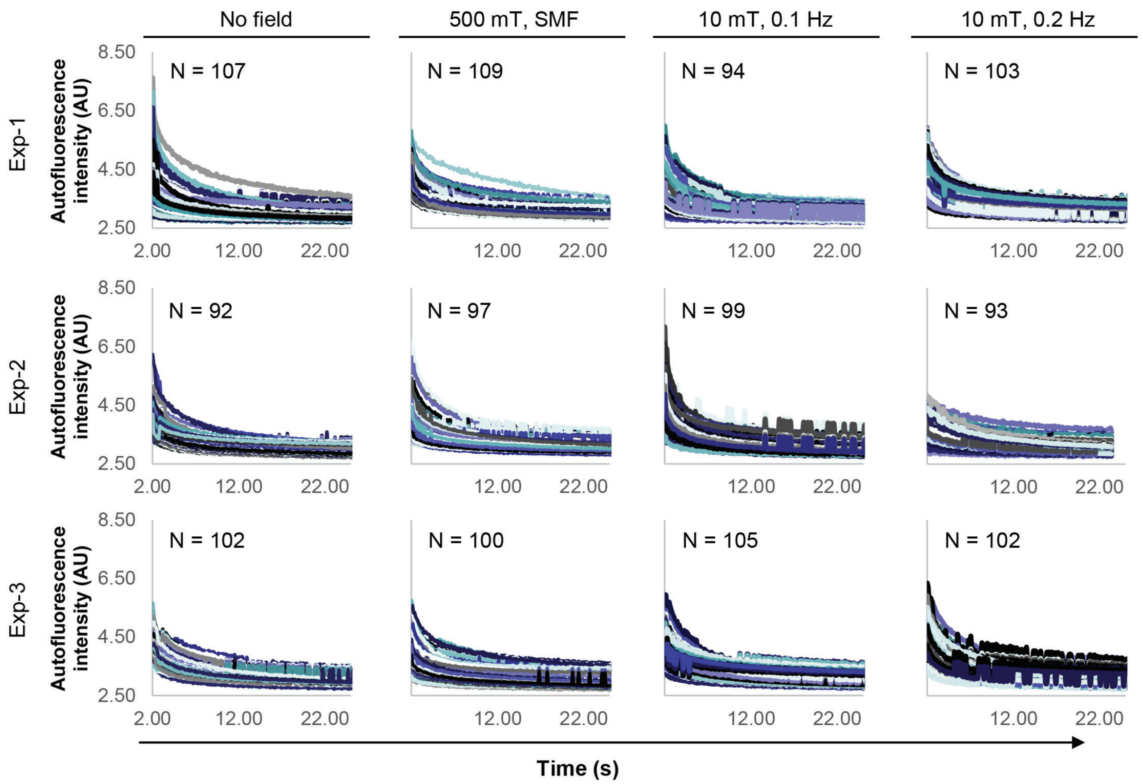


**Figure S8.** Original autoflourescence decay of HeLa cells. Cells were irradiated by 10 mT modulated magnetic field (frequencies 0.1 Hz and 0.2 Hz). 500 mT static magnetic field (SMF) was generated by bulk NdFeB magnet.

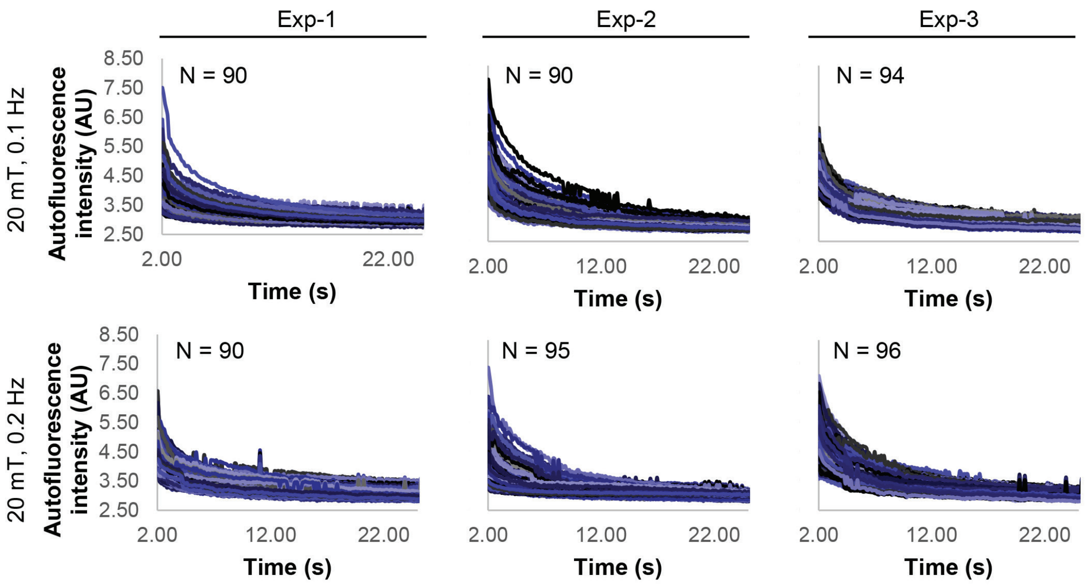


**Figure S9.** Original autoflourescence decay of HeLa cells. Cells were irradiated by 20 mT modulated magnetic field (frequencies 0.1 Hz and 0.2 Hz).

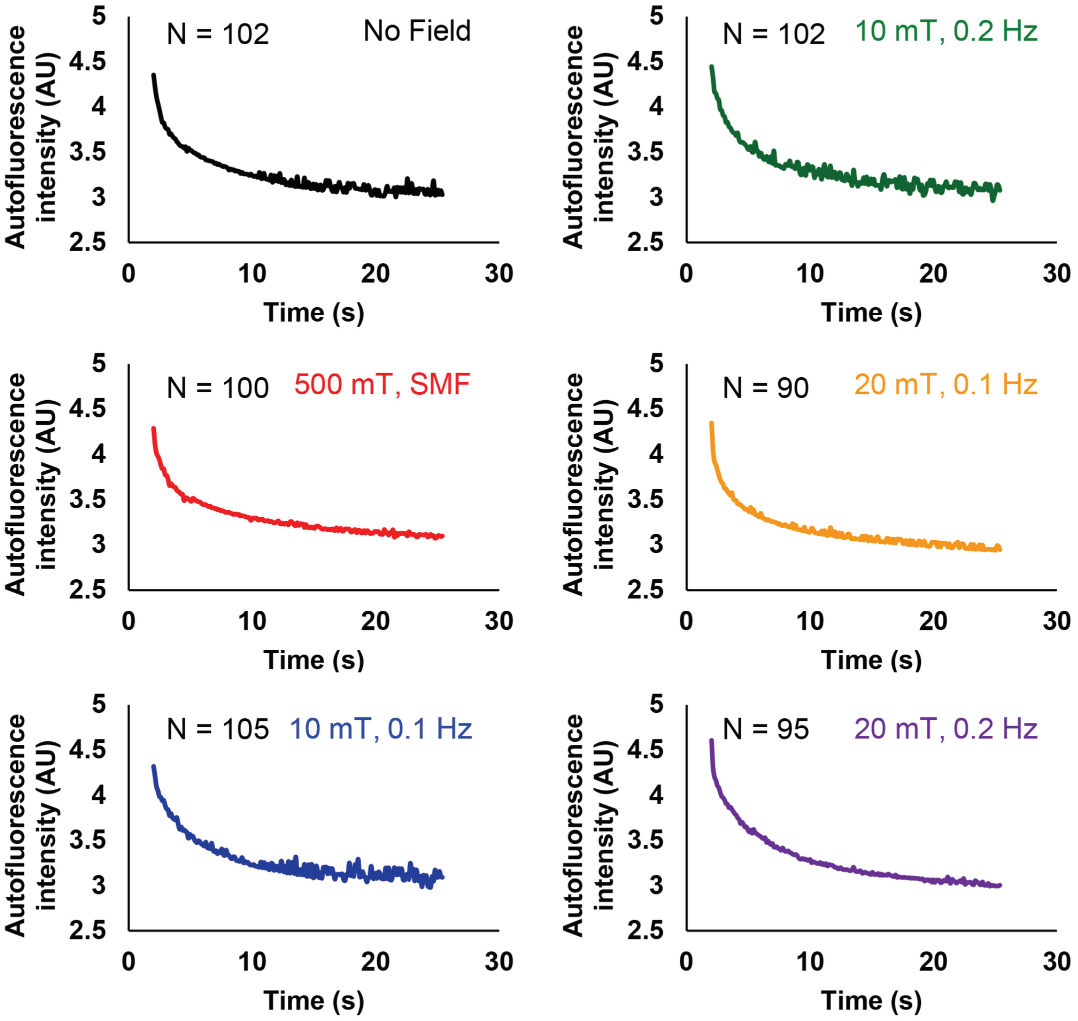


**Figure S10.** Average autofluorescence decay upon different magnetic field exposure. Cells were irradiated by 10 mT or 20 mT modulated magnetic field (frequencies 0.1 Hz and 0.2 Hz). 500 mT static magnetic field (SMF) was generated by bulk NdFeB magnet.

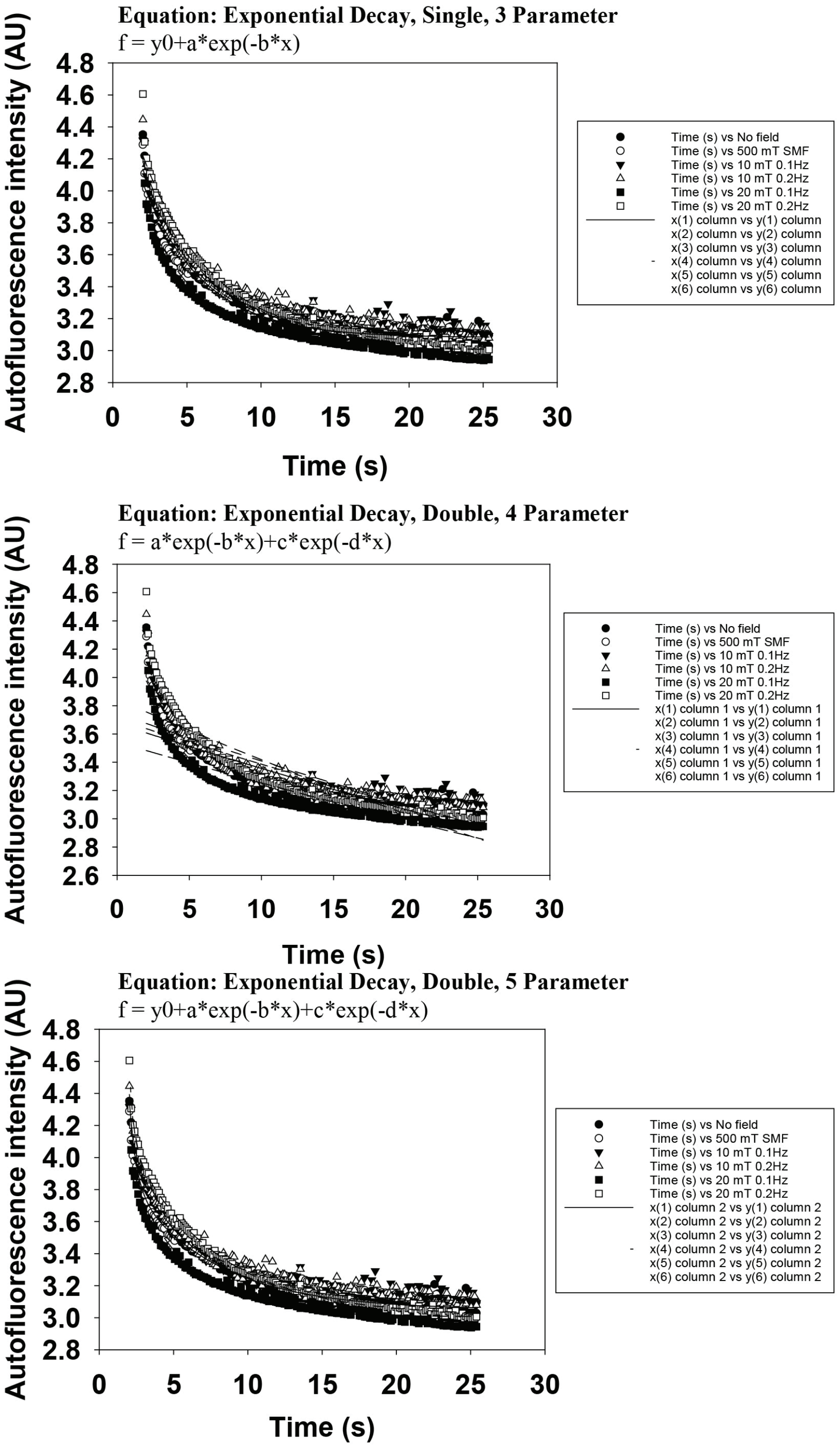


**Figure S11.** Fitting of average autofluorescent intensity decay using different functions, e.g. single exponential decay, 3 parameter; double exponential decay (Goodness of fit R = 0.9815; Rsqr = 0.9633, Adj Rsqr = 0.9627, SEE = 0.0486), 4 parameter; double exponential decay (Goodness of fit R = 0.8506; Rsqr = 0.7236, Adj Rsqr = 0.7194, SEE = 0.1305), 5 parameter (Goodness of fit R = 0.9893; Rsqr = 0.9788, Adj Rsqr = 0.9783, SEE = 0.0371). Displayed data were analyzed in SigmaPlot 13.0 software (Systat Software Inc., US).

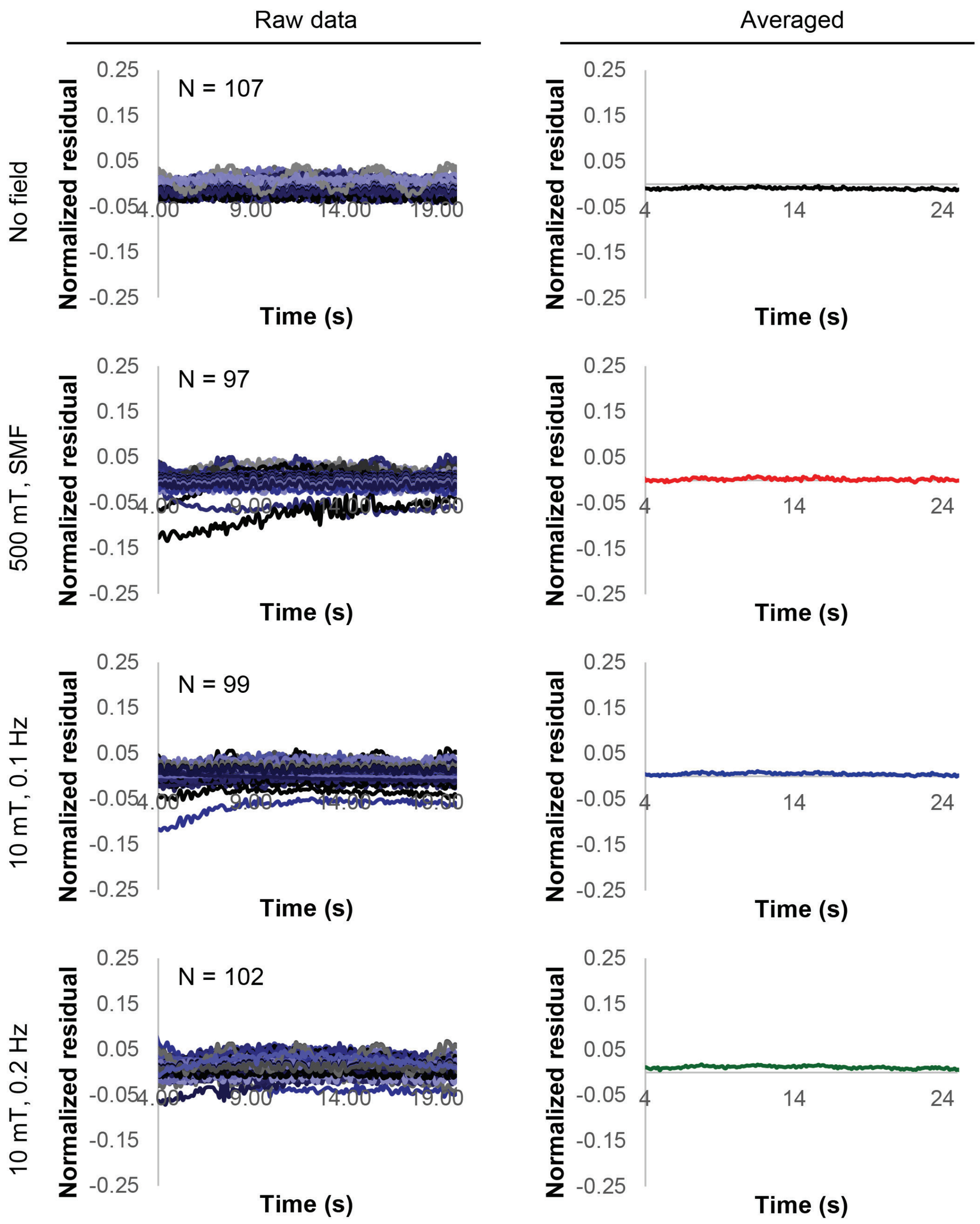


**Figure S12.** Raw data of normalized residuals of autoflourescent intensity response and corresponding averages. Cells were irradiated by 10 mT modulated magnetic field (frequencies 0.1 Hz and 0.2 Hz). 500 mT static magnetic field (SMF) was generated by bulk NdFeB magnet.

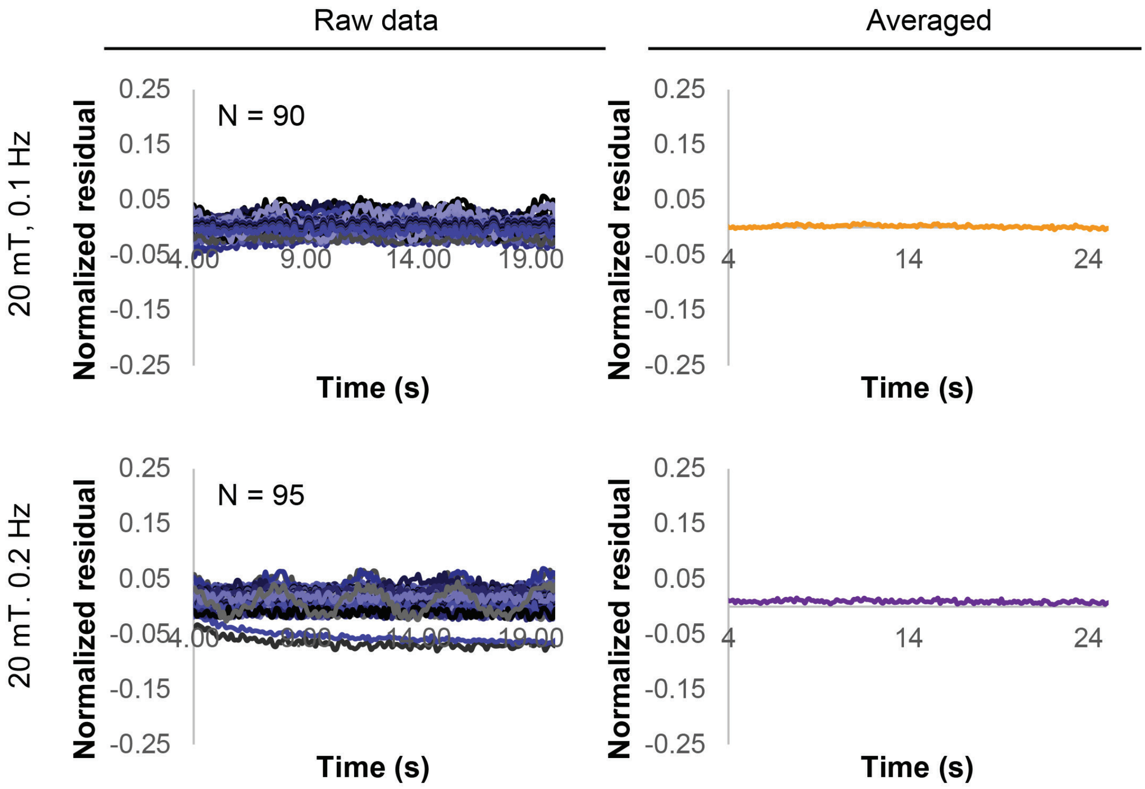


**Figure S13.** Raw data of normalized residuals of autoflourescent intensity response and corresponding averages. Cells were irradiated by 20 mT modulated magnetic field (frequencies 0.1 Hz and 0.2 Hz).

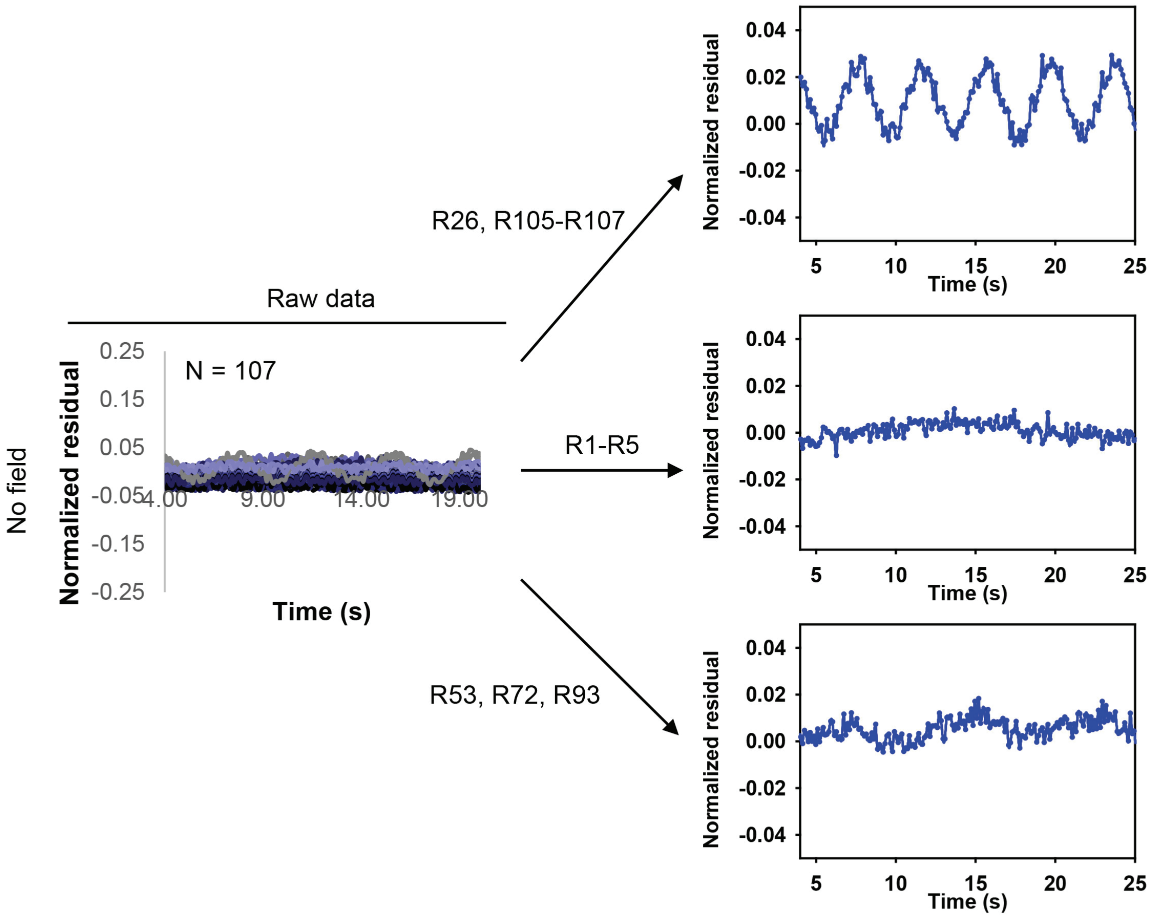


**Figure S14.** Normalized residuals of autoflourescent intensity of control cells (no field exposure). Selected cells represent different fluctuations of residuals without any magnetic field application.

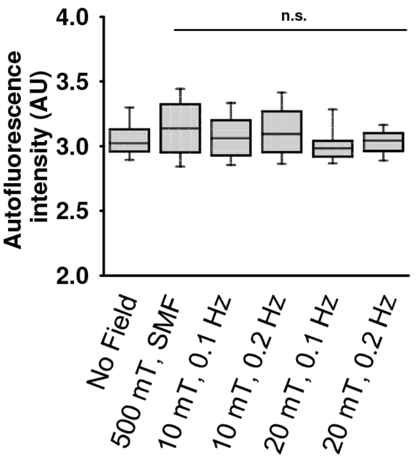


**Figure S15.** A box and whisker plot graphically represents the median, lower and upper quartiles, and lower and upper extremes of a autoflourescence intensity data. Cells were irradiated by 10 mT or 20 mT modulated magnetic field (frequencies 0.1 Hz and 0.2 Hz). 500 mT static magnetic field (SMF) was generated by bulk NdFeB magnet. N=90-107. Dunnett's test was used to determine statistical significance. Differences were considered statistically significant at $^{*}P < 0.05$.

**Legends for Movies**

**Movie S1 (separate file).** Video of a representative autofluorescence photobleaching of HeLa cells without magnetic field.

**Movie S2 (separate file).** Zoomed region of Movie S1, autofluorescence photobleaching of HeLa cells without magnetic field.

**Movie S3 (separate file).** Video showing particle tracking analysis upon autofluorescence photobleaching of HeLa cells without magnetic field.

**Movie S4 (separate file).** Video showing movable structures upon autofluorescence photobleaching of HeLa cells without magnetic field.

**Movie S5 (separate file).** Video showing non-movable and highly bleachable structures upon autofluorescence photobleaching of HeLa cells without magnetic field.

**Movie S6 (separate file).** Combined video showing particle tracking analysis upon autofluorescence photobleaching of HeLa cells without magnetic field.
